# Network analysis of patient flow in two UK acute care hospitals identifies key sub-networks for A&E performance

**DOI:** 10.1101/120188

**Authors:** Daniel M. Bean, Clive Stringer, Neeraj Beeknoo, James Teo, Richard J. B. Dobson

## Abstract

The topology of the patient flow network in a hospital is complex, comprising hundreds of overlapping patient journeys, and is a determinant of operational efficiency. To understand the network architecture of patient flow, we performed a data-driven network analysis of patient flow through two acute hospital sites of King’s College Hospital NHS Foundation Trust. Administration databases were queried for all intra-hospital patient transfers in an 18- month period and modelled as a dynamic weighted directed graph. A ‘core’ subnetwork containing only 13-17% of all edges channelled 83-90% of the patient flow, while an ‘ephemeral’ network constituted the remainder. Unsupervised cluster analysis and differential network analysis identified sub-networks where traffic is most associated with A&E performance the following day. Increased flow to clinical decision units was associated with the best A&E performance in both sites. The component analysis also detected a weekend effect on patient transfers which was not associated with performance. We have performed the first data-driven hypothesis-free analysis of patient flow which can enhance understanding of whole healthcare systems. Such analysis can drive transformation in healthcare as it has in industries such as manufacturing.

## 1 Introduction

Patient flow through a hospital is a key determinant of efficient hospital functioning. Previous studies have focused on linear flow processes, queuing and single patient pathways ^1,2^ therefore failing to capture the complex topology of a real-world hospital comprising hundreds of patient journeys overlapping in time and space.

A hospital can be considered as a network of wards that are connected when patients are transferred between them. By modelling it this way, we sought to understand both the spatial and temporal nature of network activity to help determine where and how bottlenecks form during periods of high activity. Due the complexity of the network, it was difficult to visualise network dynamics changing over time and so we took a network science approach to achieve a global, high-level perspective on hospital network dynamics.

The National Health Service (NHS) England’s Five Year Forward View proposed that IT will be an essential tool for transforming the NHS ^3^. As historically one of the most digitally mature Trusts in the UK, King’s College Hospital NHS Foundation Trust (King’s) is well-placed to use Big Data analytics to map patient flow ^4^. Using graph theory, we analysed the topology of patient flow through hospital wards at two of King’s major hospital sites modelling this as a directed flow network (Figure 1).

**Figure 1:**
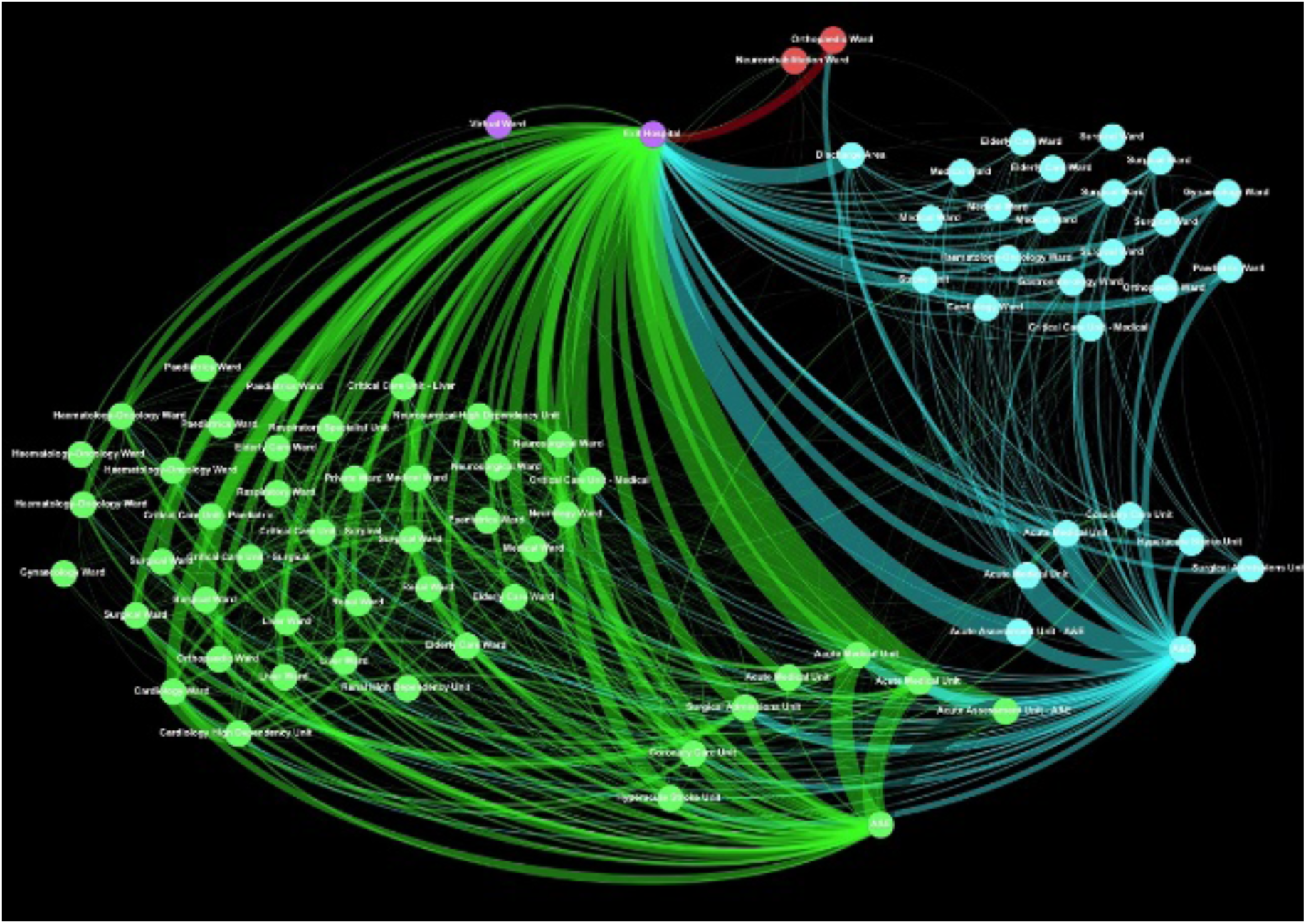
Representative topology of two of Kings’ major hospital sites over a 6-month period. Green nodes are in Denmark Hill (DH), turquoise nodes are in Princess Royal University Hospital (PRUH), purple nodes are External to King’s, red nodes are Orpington Hospital. Edge thickness represents the relative flow of the patients (see Methods below). The A&E entry points are on the lower right of the diagram. The network is directed, but for clarity at this scale direction is not represented here.

Accident and Emergency department (A&E) performance against the UK’s 4-hour waiting time target is a key metric used to assess hospitals, and patient flow through the hospital has been attributed to be a contributing factor ^5^. To understand this relationship, we performed a data-driven analysis of the major hospital sites of King’s College Hospital NHS Foundation Trust to compare network flow during periods of high performance and low performance. Although we designed our analysis to be data-driven for hypothesis-generation, we also proposed two hypotheses to evaluate the approach. We proposed that graph and cluster analysis would be able to identify (1) patterns of patient flow associated with good or poor A&E performance, and (2) cyclical patterns of patient flow on weekday and weekends.

## 2 Methods

### 2.1 Data Extraction

Data was derived from the patient administration system (PAS) and the electronic patient record at both acute sites of King’s College Hospital NHS Foundation Trust (King’s College Hospital, Denmark Hill, London [DH] and Princess Royal University Hospital [PRUH]). The PRUH-site deployed the iSoft i.PM software system (Computer Sciences Corporation, CSC) in November 2014 thereby introducing a single unified PAS across both acute sites.

A database query was performed to extract all A&E attendances and patient ward transfers between the period of 01/12/2014 and 01/07/2016. Only transfers for non-elective patients admitted via an A&E department at DH or PRUH were included.

### 2.2 Network Construction

Patient transfers were modelled as a weighted directed graph ^6,7^ in which nodes represent wards and edges represent inter-ward transfers. Edge weights are the proportion of all transfers in a given time period that each edge accounted for, allowing direct comparisons between networks with different numbers of admitted patients. Discharge from the hospital is represented by an edge to a virtual exit node. Transfers were included in the network for a site if at least one of the source or target nodes is part of that site.

### 2.3 Edge weight variability score

The edge weight variability score is a local property of a node that measures the observed deviation from a perfectly balanced distribution of its input or output edge weights as a proportion of the maximum possible deviation. If edge weights are perfectly balanced then every edge has the mean weight, so for a node n with k input edges *e*_1_, *e*_2_, …, *e*_*k*_ with edge weights *w*_1_, *w*_2_, …, *w*_*k*_, the edge weights are perfectly balanced if *w*_*i*_ =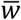 for all 1 ≤ *i* ≤ *k*. The observed deviation from the balanced situation normalised to the balanced weight is given by equation (1).

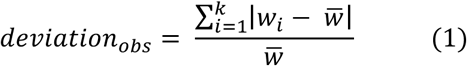

The least balanced distribution possible is the case where edges *e*_1_, *e*_2_, …, *e*_*k*−1_ have weight 1, and the remaining edge *e*_*k*_ has weight 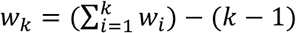 i.e. rest of the total input weight. The maximum possible deviation from the balanced state can be calculated using the above formula for the observed deviation, but substituting the vector of extreme weights for the observed weights, or equivalently by equation (2).

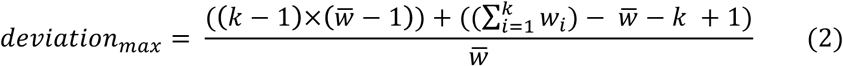

The weight variability score is now calculated by equation (3).

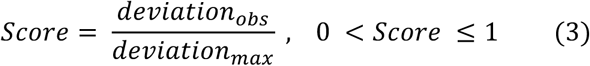

A score close to zero means the weights are balanced, a score close to 1 means the weights are unevenly distributed. This score is only defined when there are at least two (input or output) edges.

### 2.4 Cluster Analysis

Unsupervised clustering with principal component analysis was performed in R version 3.2.4 ^8^ with the weight vector for each edge centred. Edges present in less than 1% of days were removed. To determine which of the resulting components (if any) was related to performance, days were split into three categories based on A&E performance on the following day. This 1-day lag is introduced to indicate a possible causal relationship. The “high” and “low” groups were defined as the top 10% and bottom 10% of days for each site. The remaining (central) 80% of days were placed in the “middle” group.

### 2.5 Data Availability

Aggregate data are available from the corresponding author on reasonable request and subject to local information governance regulatory frameworks.

## 3 Results

### 3.1 Patient transfer networks

The patient transfers dataset contains 194,410 transfers for 78,400 hospital admissions (38,561 in PRUH, 38,115 in DH and 1,724 involving both sites) over 578 days. The weighted, directed network of all transfers analysed for the DH site contains of 78 nodes (wards) and 1531 edges, and the PRUH site has 72 nodes and 921 edges. The total number of transfers through each edge range from 1 to 11,775 in DH and 1 to 9,353 in PRUH.

From the transfer data, it is possible to reconstruct the specific path from admission to exit for each individual patient in the flow network. Many commonly used network statistics are based on the calculation of shortest paths through the network, however in this case shortest paths are not meaningful as the path a patient takes through the network is determined by the requirements of their treatment. We therefore considered the unique observed sequences of wards. Considering only those patients whose entire visit was spent in a single hospital site, there were more unique sequences of wards visited by patients in the DH site (3253) than the PRUH (2263) and the distribution of patient flow over these paths is highly uneven in both sites. Only the top 11 (DH) or 12 (PRUH) most common sequences are required to reach 50% of all visits, corresponding to 0.34% (DH) and 0.53% (PRUH) of all observed sequences of wards. In both sites the single most frequent path was an equivalent sequence from the emergency ward to a clinical decision unit and finally to exit, accounting for 21% of all visits in each case.

### 3.2 Degree stability over time

Transfer data was integrated per calendar month for each of the 19 months included in this study to capture medium-term stability of the network. The average monthly network size for DH is 919 edges (SD 60.4) and 63 nodes (SD 2.08). The PRUH networks have 592 (SD 45.4) edges and 46 (SD 2.50) nodes.

In the flow network node in-degree represents the number of other wards sending patients into a ward and out-degree represents the number of other wards patients are sent to. The difference between the in- and out-degree of a node was used as an indication of the structural role that a ward plays in the distribution of flow throughout the network and whether this changes over time (Figure 2a-b). The difference is stable over time in both sites, allowing wards to be divided into three distinct groups: in >> out, balanced, and out >> in, with almost all wards in the balanced group. Only a single ward (the clinical decision unit in DH, Figure 2b) appears to move between two categories over time, from having many more inputs to more balanced.

**Figure 2.**
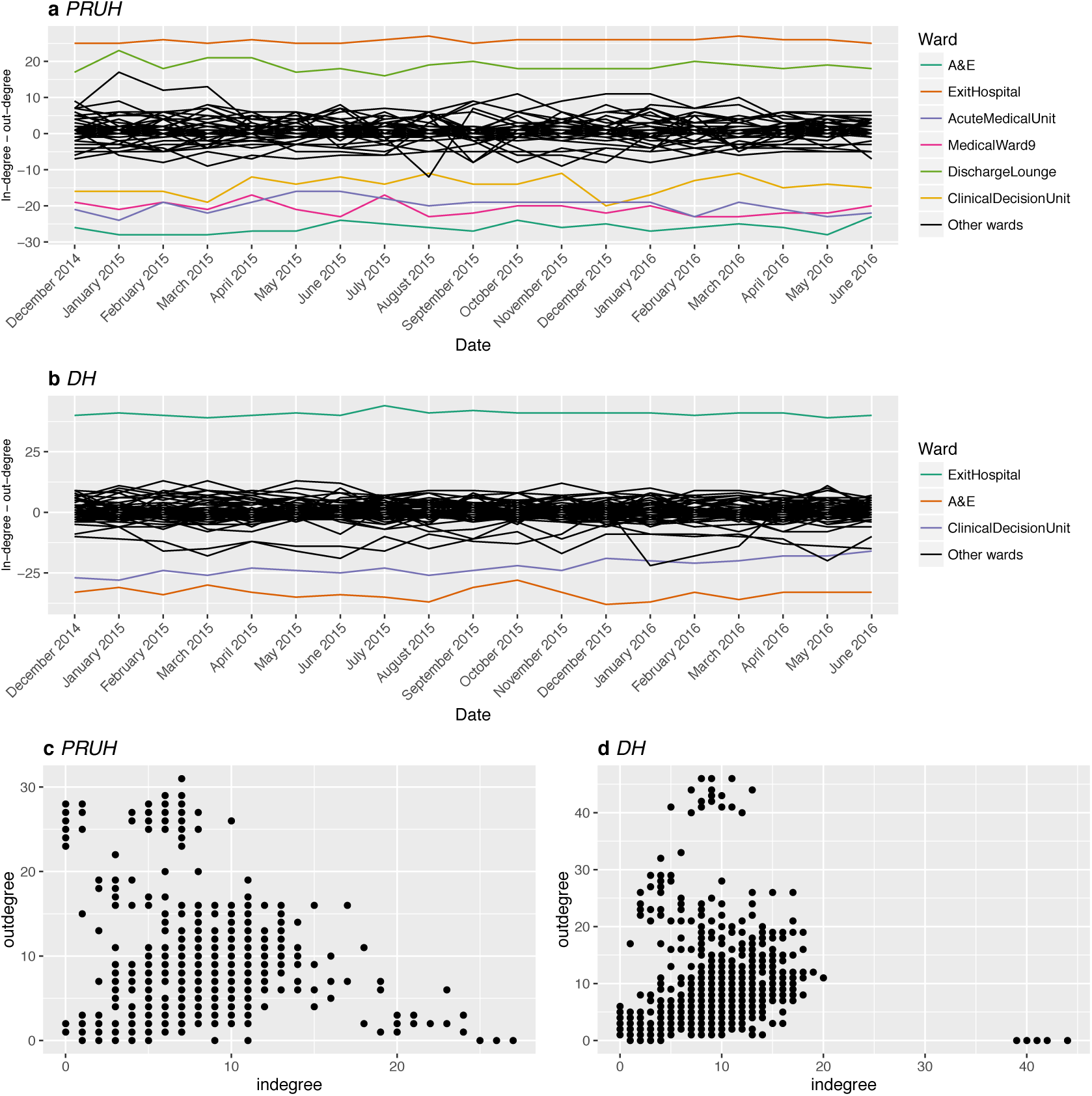
Difference between in-degree and out-degree over time. (a) and (b) Each line represents a single ward over time. Black lines represent “balanced” wards, with similar in- and out-degree. Other colours represent individual wards with either much greater in-degree than out-degree, or vice versa. Corresponding ward names for these wards are shown to the right. (c) and (d) In-degree vs. out-degree, each point is one node in one month. DH = Denmark Hill, PRUH = Princess Royal University Hospital.

From the degree difference alone, there is no way to know whether these are high or low degree nodes. Figure2c-d shows the relationship between in-degree and out-degree for every node per month. Noticeably there are no wards with both in-degree and out-degree >20 in either site.

### 3.3 Edge weight variability

The patient flow network is a weighted network. We defined a weight variability score (see Methods) that quantifies how far the input or output edge weights of a node are from being perfectly balanced (with score near 0). If the weights are as uneven as they could be, the score is 1. Applying this measure to the monthly networks showed that unlike the degree difference (Figure 2a-b) the variability score for a node is not stable over time but is consistently above 0.4 (with mean > 0.7) for both input and output weights for all nodes in both sites (Figure 3). This indicates the input and output weights are unevenly distributed, so for all nodes the majority of the input and output flow is through a small number of relatively high-weight edges. Beyond the similarity in the overall distribution of weight variability in both sites, there are specific parallels in the roles of the wards with high or low variability. In both DH and PRUH, specialist elderly/ frailty care wards have extremely uneven output edge weights and the clinical decision units and stroke units had very uneven inputs. Surprisingly, in both DH and PRUH the emergency departments themselves were amongst the most balanced in terms of output weights, with the 5^th^ (PRUH) and 6^th^ (DH) lowest output variabilities.

**Figure 3.**
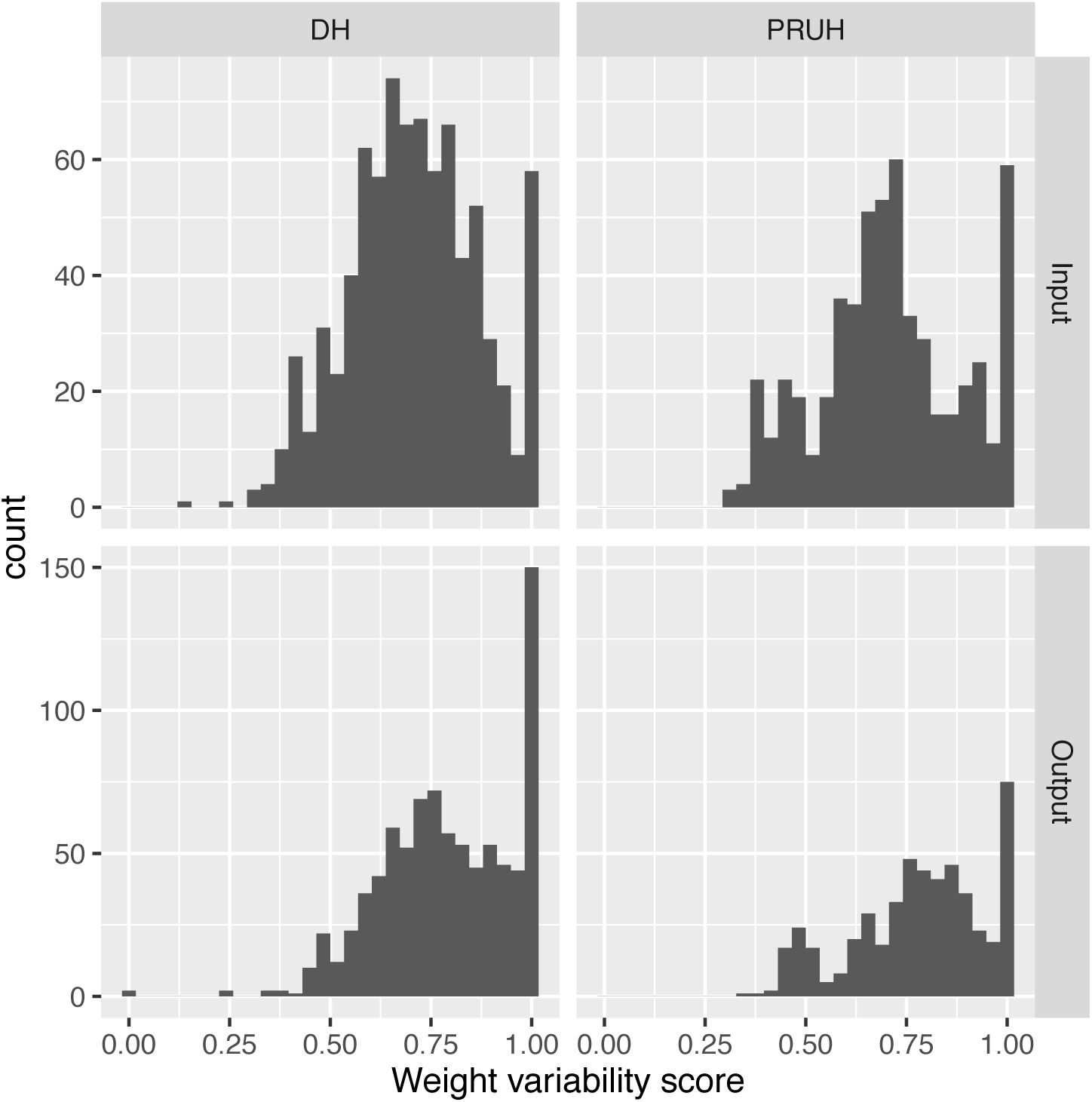
Distribution of input and output weight variability scores. The input and output weight variability scores were calculated for every node each month. Counts are over all nodes and months. DH = Denmark Hill, PRUH = Princess Royal University Hospital.

### 3.4 The “core” flow network

Edges that are consistently present over time are likely to be important to the overall functioning of a hospital, in addition to just those edges with the greatest weight. These “core” edges were defined as any edges present in all the monthly transfer networks for each site. For PRUH this leaves a subnetwork with an average of only 16.8% (SD 1.38%) of all edges per month, which accounts for 89.7% (SD 1.37%) of monthly transfers. A similar result was found for DH, where the core edges are 12.6% (SD 0.89%) of the full network on average, but support 83.4% (SD 1.54%) of all transfers. This is consistent with the above finding that that the input and output edge weights of all nodes are unevenly distributed. Therefore, although the monthly networks for both sites are quite large, a small and consistent portion of the network is sustaining the bulk of the transfer activity and is likely to be critical to maintaining patient flow.

### 3.5 Differential Network Analysis in relation to A&E performance

For the period considered in this study, the average A&E performance against the 4-hour waiting time was higher for DH than PRUH (87% vs 74.5%, Table 1), and DH is also more consistent (Figure 4). The trend over time is strikingly different between these two sites, with a clear seasonal variation in the PRUH in contrast to a recent decline in DH. The variation in A&E performance is not correlated with the number of arrivals at A&E in either site (Figure 5; linear regression R-squared = 0.14 for DH and 0.09 for PRUH). Days were divided into groups based on performance where the best and worst 10% of all days is defined as “high” and “low”, respectively, and the remaining 80% of days are in the “mid” category. Figure 4 shows the days that fall in each performance group over time. Although there is a slight tendency towards increased arrivals at A&E in the “Low” group compared to the “High” group (Table 1), the distributions are largely overlapping (Figure 5), with most days in the “Low” performance category having fewer arrivals than the maximum number in the “High” performance category (64.9% for DH and 87.7% for PRUH). Therefore, simply in terms of arrivals the network is capable of handling most of the “Low” days and performing well. A&E waiting time is thought to be partly determined by the occupancy and patient flow in downstream wards, so we considered the hypothesis that changes in the structure of the hospital network over time are correlated with changes in performance.

**Table 1.** Summary of A&E performance and arrivals per site. All values are the mean daily value with the standard deviation in brackets. “High” and “Low” groups are the top and bottom 10% of best- and worst-performing days respectively for each site, “Mid” is the remaining 80%. DH = Denmark Hill, PRUH = Princess Royal University Hospital.

|  | Performance<br>(percent meeting 4hr target) |  |  |  | Arrivals<br>(Patients) |  |  |  |
| --- | --- | --- | --- | --- | --- | --- | --- | --- |
|  | Overall<br>(n=578) | High<br>(n=57) | Mid<br>(n=464) | Low<br>(n=57) | Overall<br>(n=578) | High<br>(n=57) | Mid<br>(n=464) | Low<br>(n=57) |
| DH | 87.0<br>(6.63) | 95.5<br>(0.84) | 87.7<br>(4.50) | 73.3<br>(3.33) | 385<br>(33.2) | 366<br>(34.9) | 385<br>(32.2) | 411<br>(22.6) |
| PRUH | 74.5<br>(14.3) | 96.3<br>(1.56) | 74.9<br>(10.7) | 49.5<br>(5.91) | 177<br>(18.6) | 168<br>(15.6) | 177<br>(18.5) | 186<br>(17.8) |

**Figure 4.**
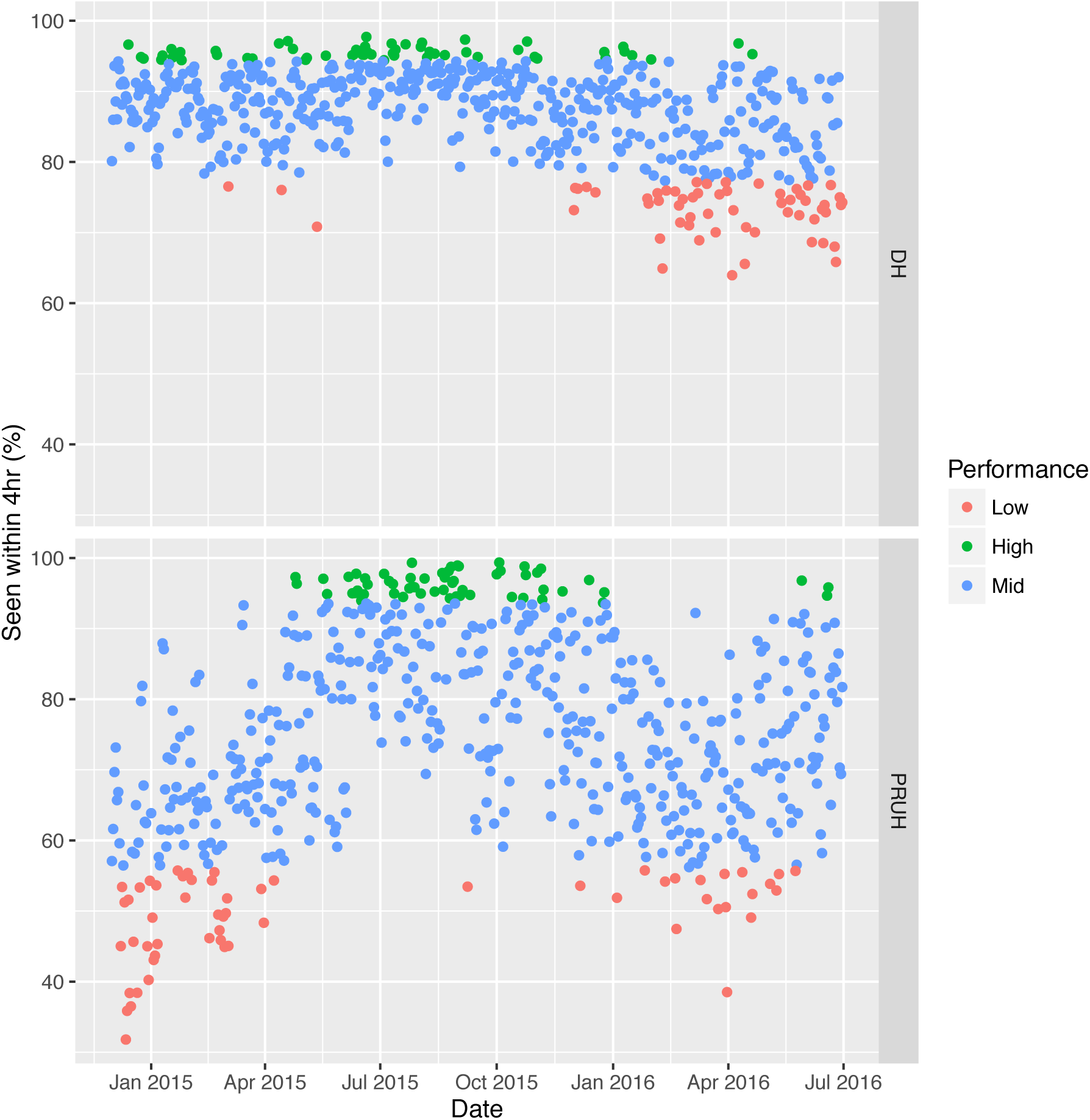
A&E performance over time in both sites. “High” and “Low” groups are the top and bottom 10% of best-and worst-performing days respectively for each site, “Mid” is the remaining 80%. DH = Denmark Hill, PRUH = Princess Royal University Hospital.

**Figure 5.**
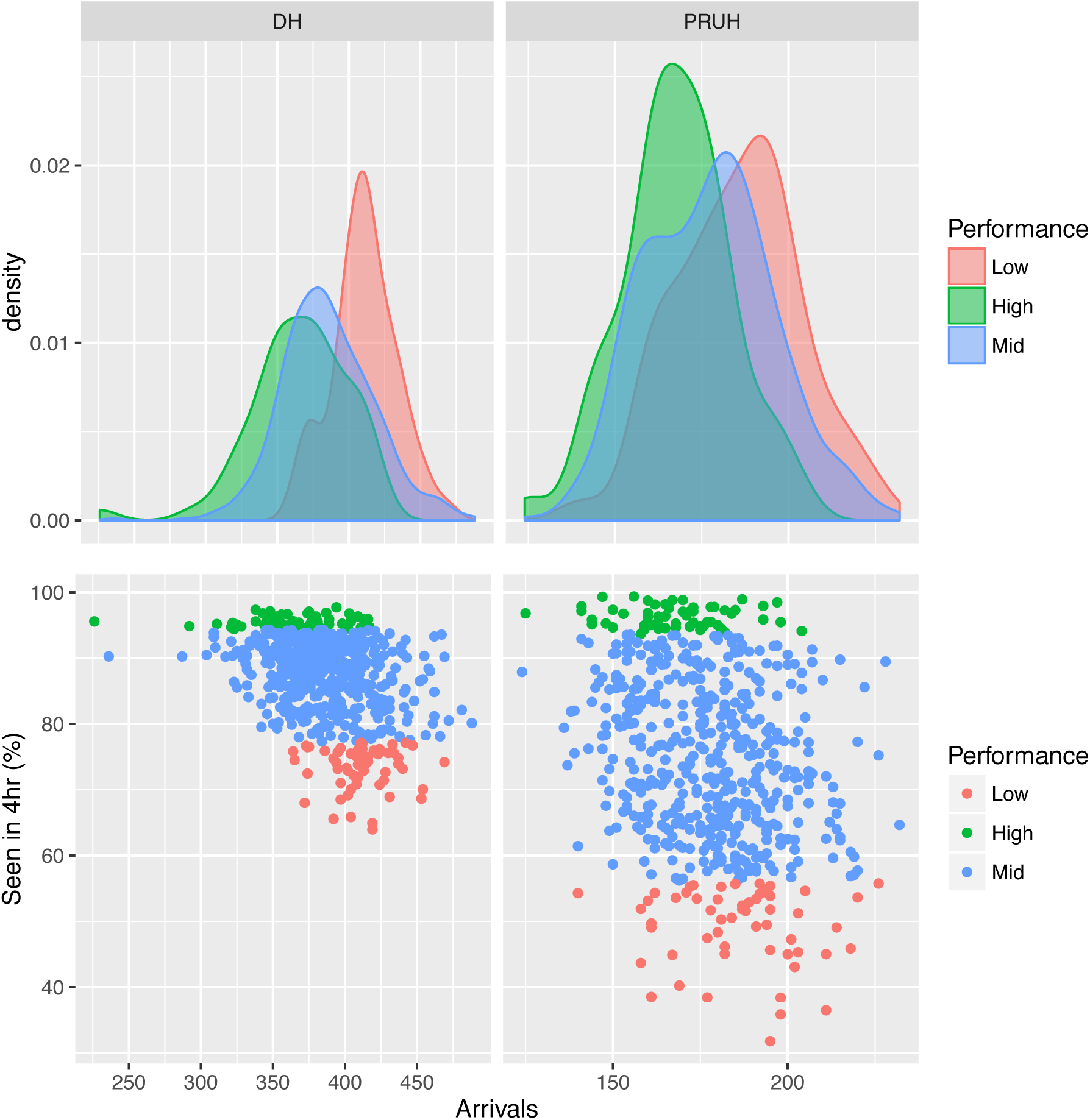
A&E performance vs. number of arrivals at A&E for both sites. The upper row shows the density of number of arrivals at A&E in each group for the points in the corresponding scatter plots in the lower row. There is no correlation between the number A&E performance against the 4-hour target and the number of arrivals at A&E (linear regression R-squared = 0.14 for DH and 0.09 for PRUH). “High” and “Low”groups are the top and bottom 10% of best- and worst-performing days respectively for each site, “Mid” is the remaining 80%. DH = Denmark Hill, PRUH = Princess Royal University Hospital.

**Figure 6.**
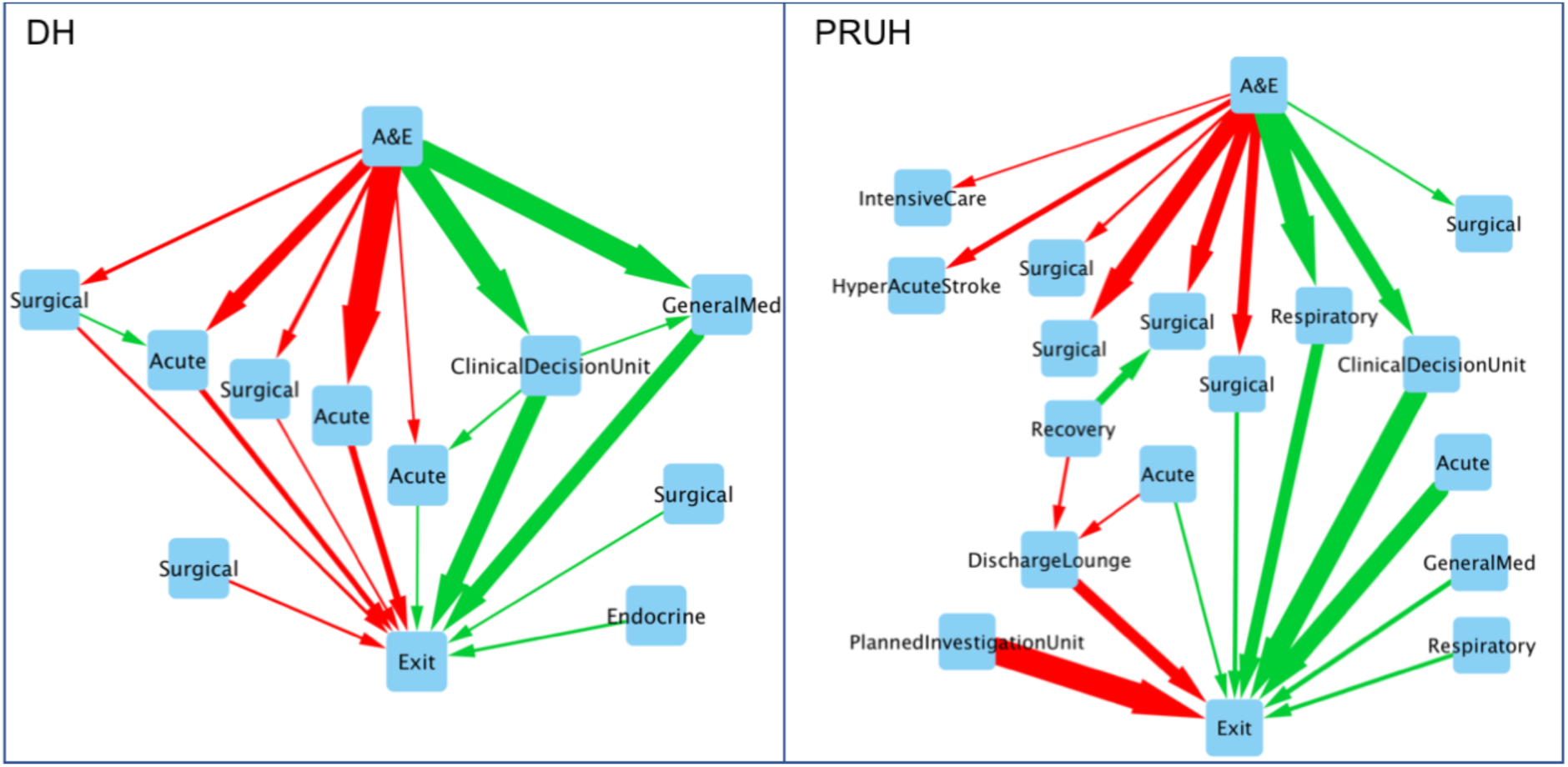
Differential networks for both sites. Only edges for which the difference in proportional flow between the best and worst networks is more than two standard deviations from the mean difference (zero) are shown in the differential networks. Red edges account for a greater proportion of transfers in the lowest-performing days, green edges account for a greater proportion of transfers in the highest-performing days. Edge direction indicates the direction of the transfer and thickness represents the magnitude of the difference in weight between the best and worst networks. DH = Denmark Hill, PRUH = Princess Royal University Hospital.

As an indication of a causal relationship (network changes driving changes in A&E performance), we looked for network changes that correlate with extreme (good or bad) A&E performance on the following day with a differential network analysis approach ^9^, focusing on edge weights. All patient transfers were pooled over the days in each group, producing a single network for each group that represents the overall structure of all the best- and worst-performing days.

Edge weights in the differential network were calculated by subtracting the weight in the worst network from the weight in the best network. An edge with a negative weight in the differential network accounts for a greater proportion of transfers in low-performing days. Weight differences between the best and worst networks were normally distributed with a mean of zero in both sites, with standard deviations of 1.32 × 10^−3^ in PRUH 2.13 × 10^−3^ in DH. Only edges with differential weights >2 standard deviations from the mean were retained in the differential network. The resulting networks are shown in Figure and represent the largest differences in patient flow on days prior to extremes of A&E performance. In DH, the edges with the largest differential weights mostly form complete paths from admission to discharge. In both sites, poor A&E performance is associated with increased flow to surgical wards the preceding day and the best A&E performance is associated with increased flow through clinical decision units, supporting their value in maintaining efficient flow through a hospital.

### 3.6 Unsupervised Clustering

Given these differences in network flow between the best and worst days, we asked whether unsupervised clustering of daily patient flow networks using principal component analysis would separate the best and worst days as defined in Section 3.5 (top 10%, bottom 10%, middle 80%). Edge weights were normalised to represent the proportion of all transfers each day to allow for comparisons between daily flow networks with different numbers of admissions. As shown in Figure 7a, the first principal component for the DH site effectively separates the high and low performance groups. The edges with the highest loadings for the first component are the same edges identified in the differential network analysis (Section 3.5).

**Figure 7.**
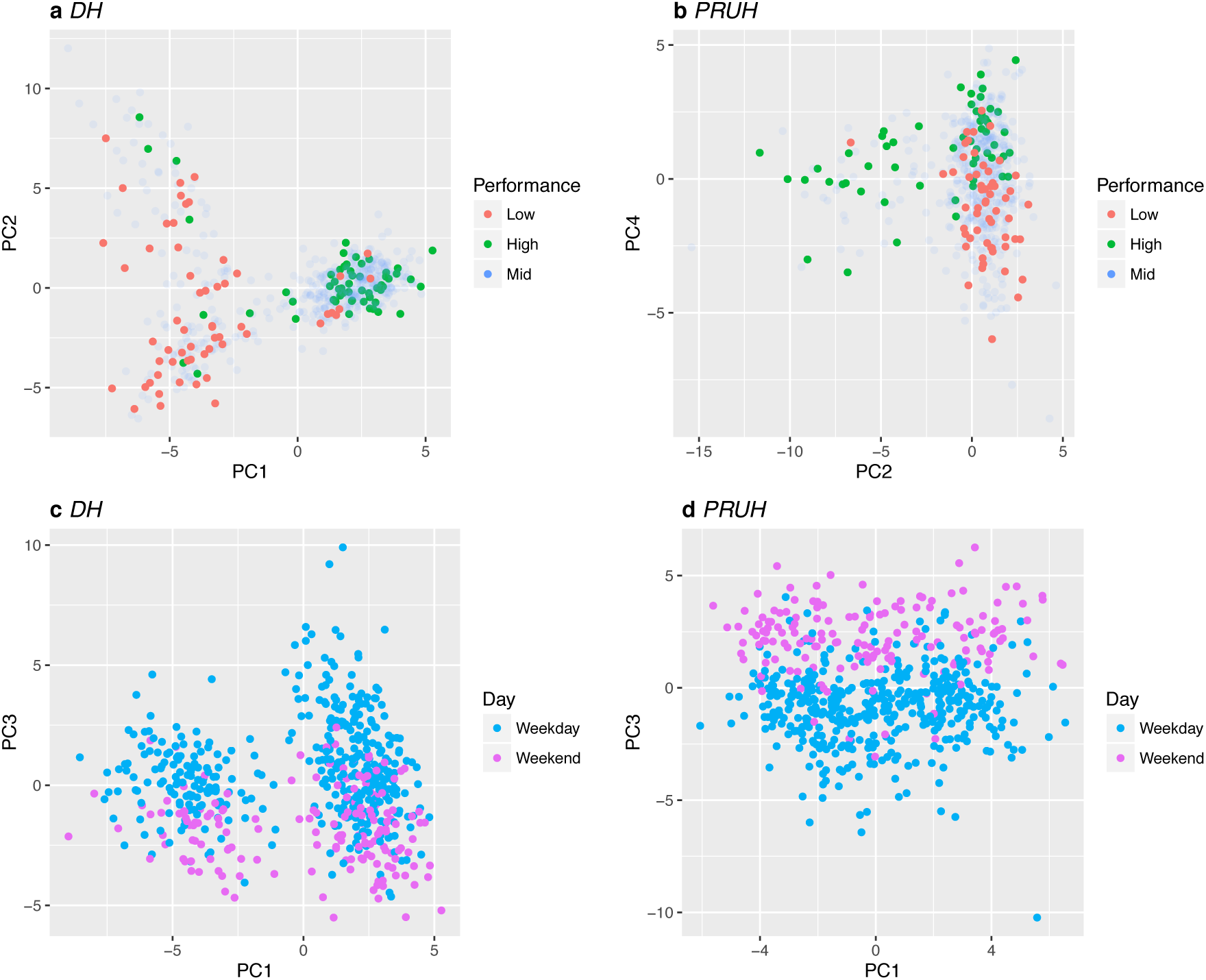
Unsupervised clustering of edge weights. Principal component analysis of daily transfer networks separates the best- and worst-performing days in each site and weekdays from weekends. (a) DH performance (b) PRUH performance (c) DH weekday vs. weekend (d) PRUH weekday vs. weekend. “High” and “Low”groups are the top and bottom 10% of best- and worst-performing days respectively for each site, “Mid” is the remaining 80%. Red = Low, blue = Mid, green = High. DH = Denmark Hill, PRUH = Princess Royal University Hospital.

The differential network analysis focused on the largest changes in edge weight, however in the PCA the weights are centred and scaled so there is no effect of magnitude. Therefore, the fact that the same edges are identified in both analyses increases our confidence that these edges are indeed related to A&E performance. For the PRUH, the separation of the best and worst days was less clear than for DH, but is still largely achieved by a combination of the 2^nd^ and 4^th^ components (Figure 7b). Again, the most important edges in these components are also identified by the differential network analysis. Overall, in both sites the differential network analysis and the PCA together implicate a consistent subset of edges from the entire flow network of each hospital as potential drivers of A&E performance. These edges are key candidates for follow-up investigation.

In both sites, the third component is correlated with whether the day is a weekday or weekend (Figure 7c-d) but does not separate days by performance. In both sites, the relative flow of patients through the clinical decision unit was one of the important variables in the third component, but with opposite patterns between the sites (less flow to the clinical decision unit on the weekend in PRUH, more flow in DH).

## 4 Discussion

We have performed a completely data-driven analysis of patient flow using modern graph theory analysis, which we believe to be the first to be done in a UK healthcare system. Our analysis shows that on a monthly basis the local network structure around wards remains constant (Figure 2). Most wards have a relatively balanced in-degree and out-degree, while very few wards act as distributors (out-degree >> in-degree). For all wards most of the input and output flow is through a small number of relatively frequent transfers, therefore each ward transfers or receives most of its patients from only a few other wards. This is consistent with our identification of a “core” sub-network in both sites that constitutes 83-90% of all flow, with the remainder flowing through atypical routes. Changes in performance in these specific areas will affect most patients, and therefore may also impact the wider hospital system. This core sub-network should therefore be the focus of any follow-up initiatives. The ability to identify key areas for in-depth review in an unbiased fashion is a key advantage of this graph theory approach to patient flow.

Most of the edges in the flow network lie outside the core sub-network and account for 10-16% of all transfers. These edges may constitute the pathways which have been least optimised. As this residual flow does not travel down the same set of edges every month, these ‘ephemeral’ sub-networks will be much harder to study. One hypothesis is that this flow constitutes less common clinical events that necessitate transfer between clinical areas made up of low-frequency ‘freak’ events (e.g. a patient having myocardial infarction after an unrelated procedure) and ‘patient safety’ events (e.g. missed opportunity to detect a deteriorating patient to prevent admission to critical care). More complex modelling will be needed to study this ephemeral sub-network

Our analysis confirmed prior data showing that A&E attendance rate is poorly correlated with hospital performance statistics ^10^. Principal component analysis and differential network analysis identified changes in the distribution of patient flow that separate the best and worst performing days in each hospital site. Flow through these pathways may be modifiable factors affecting A&E performance, however further work is required to determine whether this is a causal association, and direction of the causality. In both sites, lower A&E performance was associated with increased flow to surgical wards the previous day, but without data on individual clinical episodes we cannot determine whether this was due to a lack of beds in other wards or was required for treatment. Nonetheless, as the key factors in these components are derived from patient transfers the day before, it may be possible to incorporate them into any predictive forecasting models.

It is worth mentioning that caution must be taken when forming general hypotheses around these key pathways, as they may be specific to the local health economy for each site. One possibility is that they relate to the surrounding network of social care and community facilities that is not captured in our dataset.

Unsupervised clustering showed a weekday-weekend effect (PC3) at both hospital sites, indicating a difference in how the hospital flow network functions between weekday and weekends. This finding fits with *a priori* expectations of working patterns within the hospital and in the community. Discharge rates are lower on weekends than weekdays ^11^. So one could argue that the network changes we see are a result of working patterns of healthcare staff in hospitals and hospital operational policies.

However a large scale study showed that while changing the working patterns of senior healthcare staff (hospital consultants) did increase weekend discharges from most general medical wards, it did not have any effect on elderly medicine or geriatrics wards ^12^. Elderly medicine or geriatric units have higher proportion of individuals needing social care and the respondents of the 2015 National Inpatient Survey highlighted the bottleneck to be access to social care to allow discharge ^13^. Working patterns within the whole healthcare and social care network is likely to underlie the network differences detectable with our data-driven analysis.

It is important to be mindful that this weekend-weekday effect looks specifically at the question of patient flow and operational efficiency rather than on patient outcomes, and while the question of weekend effect on patient safety and 30-day mortality has been a topical issue in 2016 ^14–16^, it is not within the scope of this study.

In conclusion, this approach validated *a priori* hypothesis about patient flow on weekends, and generated hypotheses on what drives or impedes patient flow. It is also able to identify sub-networks within the hospital where patient flow is most greatly associated with hospital emergency department performance. It is likely that some of the factors we have identified with our analyses are specific to the local health and social care economy, but our technique can be easily exported to other health economies that capture such large volume of data.

The approach used in this study could be extended to include dynamic network analysis, extending the data capture to the entire social care economy and including other neighbouring hospital providers, and including data at the level of the single-clinical-episode within the modelling. We envisage a scenario that by pooling Big Data in such a way, it may be possible to use *in-silico* simulations for accurate forecasting within an entire healthcare economy.

## Acknowledgements

We would like to thank Jack Barker for his insightful comments on the analysis, and to express thanks to the King’s College Hospital ICT department and Electronic Health Record team for their support in this project.

This paper represents independent research funded by the National Institute for Health Research (NIHR) Biomedical Research Centre at South London and Maudsley NHS Foundation Trust and King’s College London. The views expressed are those of the author(s) and not necessarily those of the NHS, the NIHR or the Department of Health. This research was supported by researchers at the National Institute for Health Research University College London Hospitals Biomedical Research Centre, and by awards establishing the Farr Institute of Health Informatics Research at UCL Partners, from the Medical Research Council, Arthritis Research UK, British Heart Foundation, Cancer Research UK, Chief Scientist Office, Economic and Social Research Council, Engineering and Physical Sciences Research Council, National Institute for Health Research, National Institute for Social Care and Health Research, and Wellcome Trust (grant MR/K006584/1).

## Author contributions

J.T., D.M.B and R.J.B.D. designed the study; J.T. collected the data; D.M.B carried out analysis; J.T., R.J.B.D, N.B and C.S. advised on analysis; D.M.B, J.T. and R.J.B.D prepared the manuscript; all authors revised the manuscript.

